# Effective networks mediate right hemispheric dominance of human 40 Hz auditory steady-state response

**DOI:** 10.1101/2023.02.02.526849

**Authors:** Neeraj Kumar, Amit Jaiswal, Dipanjan Roy, Arpan Banerjee

## Abstract

Auditory steady-state responses (ASSR) are induced from the brainstem to the neocortex when humans hear periodic amplitude-modulated tonal signals. ASSRs have been argued to be a key marker of auditory temporal processing and pathological reorganization of ASSR – a biomarker of neurodegenerative disorders. However, most of the earlier studies reporting the neural basis of ASSRs were focused on looking at individual brain regions. Here, we seek to characterize the large-scale directed information flow among cortical sources of ASSR entrained by 40 Hz external signals. Entrained brain rhythms with power peaking at 40 Hz were generated using both monaural and binaural tonal stimulation. First, we confirm the presence of ASSRs and their well-known right hemispheric dominance during binaural and both monaural conditions. Thereafter, reconstruction of source activity employing individual anatomy of the participant and subsequent network analysis revealed that while the sources are common among different stimulation conditions, differential levels of source activation and differential patterns of directed information flow using Granger causality among sources underlie processing of binaurally and monaurally presented tones. Particularly, we show bidirectional interactions involving the right superior temporal gyrus and inferior frontal gyrus underlie right hemispheric dominance of 40 Hz ASSR during both monaural and binaural conditions. On the other hand, for monaural conditions, the strength of inter-hemispheric flow from left primary auditory areas to right superior temporal areas followed a pattern that comply with the generally observed contralateral dominance of sensory signal processing.

## 1. Introduction

Auditory steady-state response (ASSR) is a phase-locked oscillatory response to periodic sound recorded from the scalp (Picton 2013; Zhang et al., 2013). ASSRs can be identified by the frequency-based analysis, exhibiting a sharp rise in spectral power and phase-locking across trials at the frequency of periodicity. In the literature, ASSRs have been reported widely to be elicited maximally at a stimulation frequency of 40 Hz (Galambos et al., 1981; Hari et al., 1989; Pastor et al., 2002). Moreover, ASSRs can be detected readily at a single participant level and do not show gender-specific differences among right-handed participants (McFadden et al., 2014; Melynyte et al., 2017). Due to its robustness and reliability, 40 Hz ASSR is widely used for theoretical and clinical research, e.g., temporal auditory processing, screening hearing threshold, and measurement of consciousness during global anaesthesia (Farahani et al., 2020; Haghighi et al., 2018; Niepel et al., 2020), etc. Additionally, 40 Hz ASSR is also used as a biomarker in certain neuropsychological disorders like schizophrenia and autism (O’Donnell et al., 2013).

In general, the processing of sensory stimuli requires coordinated interactions among specialised brain regions that are distributed across the hierarchal cortical networks (Bressler & Menon, 2010; Fries, 2005). Lithari and colleagues have reported the emergence of a frequency-specific large-scale network during the visual steady-state response, indicating synchronized activity across multiple regions of the brain in response to visual stimulation (Lithari et al., 2016). Therefore, studying the pattern of information flow among specialised brain regions that are involved would allow us to understand the neural basis of ASSRs. There is converging evidence that neural generators for 40 Hz ASSRs predominantly lie in the right superior temporal gyri (STG) (Edgar et al., 2016; Kim et al., 2019) and bilateral primary auditory cortices (Bohórquez and Özdamar 2008; Pantev et al., 1996; Ross et al., 2005; Steinmann et al., 2011) in addition to sub-cortical regions (Herdman et al., 2002; Poelmans et al., 2012; Steinmann et al., 2011). Despite the growing consensus on the regions involved in the activation of 40 Hz ASSRs, the interactions and information flow among the relevant nodes of ASSRs are yet to be fully investigated. Importantly, the patterns of information flow between relevant brain regions in the distributed auditory hierarchy are indicative of directed functional connectivity, which offers a refined picture of the communication channels and a critical role of the underlying drivers and followers involved in the generation of ASSRs (Bressler & Menon, 2010; Lithari et al., 2016). For example, a unidirectional information flow from one sensory node to a higher-order will reveal hierarchical processing. On the other hand, a bidirectional connectivity between cortical nodes would imply a parallel and simultaneously recurrent processing scheme. For instance, the flow of information between left and right auditory cortical nodes are mediated via strong cortico-thalamocortical feedback loops (Das et al., 2021; Li et al., 2018). Thus, an overall characterization of the direction of information flow pathways in auditory pathways within and between two hemispheres will reveal that the functional organization between two hemispheres is governed by the symmetry/ asymmetry of auditory stimulation during auditory tone processing tasks. Motivating our work from previous studies that suggest lateralization enables more efficient information transfer among specialized brain regions during language processing (Toga et al. 2003), we propose studying ASSR lateralization at the cortical source space level and interactions between key cortical sources. We hypothesize that causal interactions between cortical sources will provide critical insights into underlying brain networks and their task-specific functional organization. Following studies demonstrating right hemispheric dominance of non-linguistic / tonal/musical auditory sounds (Albouy et al., 2020), we further hypothesize that the underlying effective brain network interactions (Friston, 2011) should also reflect the presence of hemispheric dominance. Furthermore, the directionality of effective connections will reflect the hierarchical aspects of underlying information processing.

Earlier studies have reported right hemispheric dominance in source activation during 40 Hz ASSRs. Existing evidence from structural and functional studies suggests that ~70 % of ascending auditory inputs from either ear cross at the brainstem level and 30% remain on the same side (Hackett, 2015; Kaas & Hackett, 2000; Langers et al., 2005). Consequently, one would expect a contralateral dominance in primary auditory cortical (PAC) response during monaural stimulation conditions. Now, it’s naturally intriguing how the information, particularly during the monaural right condition, that enters the left PAC is eventually redistributed to a specialised centre present in the right hemisphere. Transcortical communication through the corpus callosum (CC) has been suggested to compensate for this asymmetric input to achieve hemispheric specialization during the processing of distinct features of auditory stimuli (Zaidel & Iacoboni, 2003; Cammoun et al., 2015). This can be very well experimentally validated by monaural stimulations. Hence, according to the principle of contralateral dominance, the monaural right would evoke a greater response in the left PAC than the monaural left condition (Andoh, Matsushita, and Zatorre 2015). In the present work, we record human electroencephalography (EEG) during 40 Hz ASSRs during binaural and both monaural left and right stimulation conditions. We reconstruct trial-wise source activity employing subject-wise anatomical structure by co-registering EEG with individual subject Magnetic resonance imaging (MRI) data. We first confirm the presence of ASSR and its well-known right hemispheric dominance co-existing with contralateral dominance of early auditory processing. Subsequently, we characterize effective network interactions using spectral Granger’s causality among sources of ASSRs.

## 2. Materials and Methods

### 2.1. Participants

Twenty-one healthy, right-handed human volunteers (16 males, 5 females, age range 22–39 years old; mean ± SD = 28 ± 2.10) participated in this study^1^. The right-handedness of participants was verified by the Edinburgh Handedness Questionnaire based upon a cut-off of 60–100. All the volunteers reported no medical history of audiological, neurological or psychiatric disorders. All of them had normal or corrected to normal visual acuity. Informed consent was given by all the volunteers in a format approved by the Institutional Human Ethics Committee (IHEC) of National Brain Research Centre that confirms the guidelines set by the Declaration at Helsinki. All participants were fluent in at least two languages, Hindi and English, but some were familiar with another language of Indian origin.

### 2.2. Experimental design

Stimuli consisted of sinusoidal tones of frequency 1 kHz and 25 ms duration, presented 40 times per second. Wherein, each tone of 25 ms had 2 ms fade in and faded out period (Figure 1A: Upper panel). Each trial comprised of 1s “On” block (auditory stimulation) period followed by 1s “Off” block (silent) period (Figure 1A). A total of 100 trials were presented for each kind of auditory stimulation, monaural and binaural. In total, four experimental conditions, each lasting 200 seconds, were performed in the following order— 1) a baseline condition in which the volunteers were not given any tonal stimuli; 2) Binaural (in both ears); 3) Monaural left (only through left ear); 4) Monaural right (only through right ear). The time interval between each condition was set to 100 s (silent). Auditory stimuli were created and presented in Stim2 software (Compumedics, Inc., USA) at 85 dB sound pressure level. The specific value of intensity was chosen to have reliable ASSR responses in all participants based on a prior pilot study in our group. Participants were instructed to stay still in a sitting position, fixate on a visual cross displayed on a computer screen for the duration, and listen to the tones. Continuous scalp EEG was recorded when the volunteers were performing the experiment.

**Figure 1.**
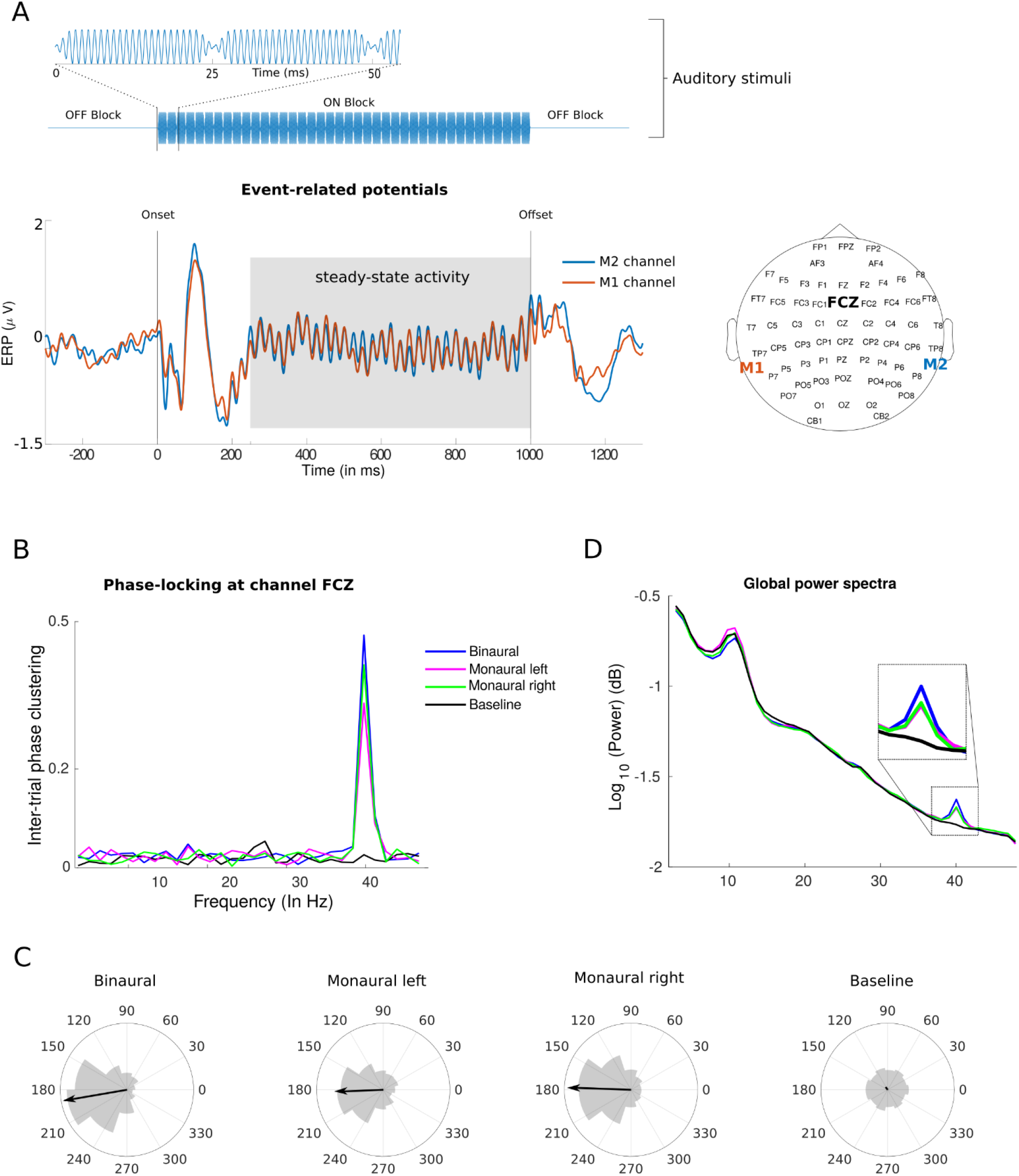
Steady-state response: (A) Stimuli (Upper panel): 25 ms of pure tone (1 kHz frequency), presented 40 times in a second during 1 s ON block interspersed by two OFF blocks (silent). Lower panel: Group-level ERPs of mastoid channels (M1 as orange and M2 as sky-blue) having oscillatory response in the time window of 250 to 1000 ms relative to the onset of periodic auditory stimuli. (B) ITPC of single channel (D) Group-level power spectrum showing sharp enhancement at 40 Hz during monaural left (magenta), monaural right (green), binaural stimuli (blue) relative to baseline (black) condition. (C) Phase-angle distribution at 40 Hz during auditory stimulation and baseline conditions. The arrow in the middle of each circle represents strength of ITPC.

### 2.3. Data acquisition

EEG data were recorded using 64 Ag/AgCl sintered electrodes mounted in an elastic head cap according to the international 10-20 system. All recordings were done in a noise-proof isolated room using NeuroScan (SynAmps2) system (Compumedics Inc, USA) with a 1 kHz sampling rate. Abrasive electrolyte gel (EASYCAP) was used to make contact between EEG sensors and scalp surface, and impedance was kept below 5 kΩ in each sensor. The default EEG system-assigned reference electrode was placed at the vertex (Cz) and the ground electrode at the forehead (AFz). The electrode locations were obtained relative to three fiducials at the nasion and left and right preauricular points using a 3D digitizer (Polhemus Inc., Colchester, VT, USA).

### 2.4. Pre-processing of EEG signals

EEG data analysis was performed with Chronux (http://chronux.org/), EEGLAB (Delorme & Makeig, 2004), FieldTrip (http://fieldtriptoolbox.org) and custom MATLAB scripts (www.mathworks.com). EEG data was imported in MATLAB using EEGLAB from Neuroscan raw files. Continuous long-time EEG time series were bandpass filtered to retain frequencies between 3–48 Hz. Thereafter, eyeblinks and heartbeat artefacts were removed from EEG data using independent component analysis in-built in EEGLAB after careful visual inspection. The temporal window of steady-state activity was identified from the grand mean event-related potentials (ERPs) of both mastoid channels (Coffey et al., 2016) (Figure 1). Hence, baseline corrected epochs of 750 ms from “On” blocks were extracted from each trial, excluding the first 250 ms of the time series from the onset of auditory stimuli. Furthermore, simple threshold-based artefact rejection was applied to exclude the remaining noisy epochs. Hence, epochs having a voltage greater than ±85 μv were discarded. After artefact rejection, none of the epochs from monaural conditions of one participant survived the thresholding. Therefore, we removed the data of that participant from further analysis. Thereafter, we re-referenced the EEG signal to common mode average reference, followed by combining trials from every participant into a single pool for group analysis at the sensor-level. On average less than 5 trials were removed per subject. In total we have recorded 2000 trials from all participants per condition (100 trials X 20 participants). The surviving trials numbers were 1980 in binaural; 1975 for monaural left and 1980 trials for monaural right.

### 2.5. Identification of ASSRs at sensor-level

#### 2.5.1. Inter-trial phase clustering

Inter-trial phase clustering (ITPC) was employed to ensure the presence of the phase-locked oscillatory response of 40 Hz. ITPC quantify the degree of nonuniformity or clustering in the distribution of oscillatory phase across trials at a particular frequency (Palva et al., 2005). The value of ITPC range between zero and one, with one being perfect phase consistency across trials and zero being complete uniformity. Mathematically, ITPC is defined as

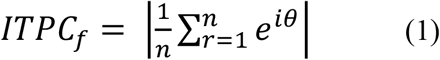

wherein, 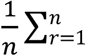 represents an average of complex vector *e^iθ^* across *n* trials; *e^iθ^* is Euler’s formula wherein *θ* is phase angle at frequency *f* derived from complex Fourier coefficients. We calculated the ITPC of all the channels during the binaural condition and selected a channel with maximum ITPC at 40 Hz. To assess the significance of phase clustering, the observed ITPC values were compared with a critical value *ITPC_crit_* computed at the significance level of p = 0.01 (Bonferroni corrected). The critical value corresponding to *p*-value was obtained as (Zar, 1999, Cohen 2014)

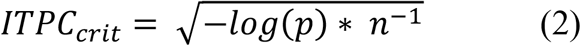

where n is the total number of trials. ITPC values higher than *ITPC_crit_* were considered statistically significant.

#### 2.5.2. Power spectrum

The power spectrum was computed using Chronux function *mtspectrumc.m* scripts at each sensor, trial, and condition. First, power spectra were calculated in the frequency range of 3–48 Hz (~ 1 Hz smoothing), and grand averaged for all channels and trials. Subsequently, t-statistic were calculated as test statistics between auditory stimulation and baseline conditions at the frequency of interest i.e., 40 Hz, followed by a statistical evaluation of observed t-statistic employing a permutation test described above. The procedure involves 1000 time shuffling of the trials among both conditions and measuring t-statistic in each permuted data sets. Thereafter, a histogram is plotted, comprising all 1000 t-statistic from permuted data creating a null distribution (Maris & Oostenveld, 2007). Observed t-statistic were then thresholded to the 99th quantile of the respective null distribution corresponding to p = 0.01.

#### 2.5.3. Laterality analysis

Hemispheric asymmetry in brain responses was quantified using laterality index (*LI*), which is the difference between right hemisphere (RH) and left hemispheric (LH) responses normalized by the sum of responses in both hemispheres.

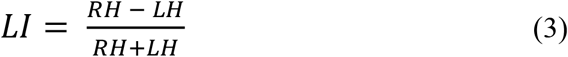

The value of *LI* ranges between +1 and −1. Wherein +1 represents complete right hemispheric dominance, −1 for complete left hemispheric dominance and 0 for a bilaterally symmetric response. Trial-wise median of 40 Hz spectral power and ITPC was calculated over right and left hemispheric sensors, excluding midline sagittal plane electrodes. Furthermore, to assess the statistical significance of LIs, the 95% confidence interval (*CI*) was calculated as

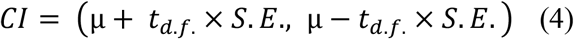

where μ is the mean of data, *t* is the inverse of Student’s t cumulative distribution function at corresponding *d. f*(degree of freedom), and *S. E*. is the standard error of the mean. Moreover, to examine whether lateralization indices of different auditory conditions have equal means, we performed one-way ANOVA on the distribution of LI values of different stimulation conditions, followed by controlling for multiple comparisons employing the Tukey-Kramer post-hoc test.

### 2.6. Source localization of ASSRs

Converging evidence suggests that utilising individual brain anatomy yields better source localization (Coffey et al., 2016). Hence, source-level analysis was performed by co-registering EEG sensor locations to MRI-guided fiducial points. Exact low-resolution brain electromagnetic tomography (eLORETA) (Pascual-Marqui, 2007) was used to calculate the three-dimensional spatial distribution of source activity underlying 40 Hz ASSRs. Earlier research demonstrated that eLORETA yields a favourable performance when false positives were considered (Halder et al., 2019). eLORETA employs distributed source modelling and estimates the current source density across brain volume by minimizing the surface Laplacian component during the construction of the spatial filter (Pascual-Marqui, 2007). Additionally, eLORETA does not rely upon any assumption regarding the number of underlying sources while having excellent control over the suppression of false positives during the detection of sources (Halder et al., 2019). Hence, source analysis employing eLORETA was performed using the FieldTrip toolbox (Oostenveld et al., 2011) implemented in MATLAB and connectivity analysis was done by customized MATLAB scripts.

The ingredients to construct a frequency domain eLORETA spatial filter are the forward model and the cross-spectral matrix of sensor data. The forward model also called the volume conduction model for the head define how an electrical current propagates across brain and would be recorded at the sensor level. Therefore, geometrical properties of the head including surface description of brain, skull and scalp were derived from subject-specific T1-weighted anatomical MRI scan and used for forward modelling based on the boundary element method (Fuchs et al., 2001). Subsequently, employing channel position leadfields for individual subjects were computed. Construction of leadfield requires description of locations known as grids on which the leadfields are calculated. Here, we first create 11,000 template grids that were defined according to Automated anatomical labelling (AAL) atlas (Rolls et al., 2015). Subsequently, these template grids were warped to individual MRI yielding 11,000 subject-specific grids arranged in normalised space. The leadfield matrix was computed at each grid in 3 orthogonal directions.

Moreover, we computed sensor-level cross-spectral matrices for each condition from the SSR time series (250:1000 ms), same as used for the sensor-level analysis (Figure 1A). Thus, we computed a spatial filter employing the subject-specific forward model and sensor-level cross-spectral matrix for each condition. A common spatial filter was computed from combined data i.e., trials from all four conditions were grouped into a single pool. Since, common filter employs cross-spectral matrices from all conditions hence, attenuating filter-specific variability during inverse modelling, i.e., the observed difference between different conditions is attributed only to the differences in conditions, not due to differences in the spatial filter. Subsequently, sensor-level cross-spectra were projected to the spatial filter obtaining the source power across trials, space and orientation. Since we do not have any prior assumption about the orientation of the underlying source activity, the largest eigenvalue was selected as grid power from strongest dipole orientation that corresponds to the maximum variance of data. Consequently, subject and trial wise distribution of source power across brain volume was obtained for each condition. Thereafter, pairwise t-statistic was computed as test statistic for each grid between auditory stimulation and baseline condition. Since there was total 11,000 grids hence would have required same number of statistical comparisons, therefore, to circumvent the multiple comparison problem we down-sampled the source powers from grid space to parcel space by taking median t-statistic of each parcel. Further, the prominent sources for each subject were selected after thresholding at the 95^th^ quantile from distribution of t-statistic of parcels and further tested for statistical significance by non-parametric statistic as described above for sensor-level 40 Hz spectral power (see section 2.5.2). Finally, for visualization purpose, all grids from significant parcels were visualized after rendering onto a cortical surface from the Colin27 brain template provided in the FieldTrip toolbox (http://fieldtriptoolbox.org).

### 2.7. Source activity reconstruction and connectivity analysis

The source-level Fourier coefficients were reconstructed by projecting the trial-wise sensor-level cross-spectral matrix to the spatial filter of significant parcels. These spectral coefficients were utilized to calculate global coherence, ITPC (see supplementary information) and Granger’s causality among sources.

#### 2.7.1. Global Coherence

Global coherence analysis was employed to identify the presence of a brain-wide large-scale functional network (Cimenser et al., 2011, Fonseca et al., 2015). The global coherence can be calculated from the cross-spectral matrix using the leading eigenvalue method in two steps. First, cross-spectrum was computed as:

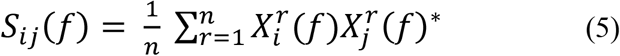

where 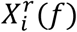 and 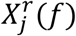 are trial (*r*) and frequency (f) specific Fourier coefficients from the sources *i* and *j*, respectively and asterisk represents matrix transposition and complex conjugate calculated over *n* trials. Second, global coherence was computed as the ratio of the maximum eigenvalue and the sum of eigenvalues calculated from the cross-spectral matrices.

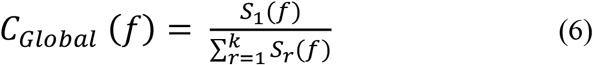

where *C_Global_*(*f*) is the global coherence, *S*_1_(*f*) is the largest eigenvalue and the denominator 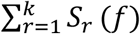 represents the sum of eigenvalues of the cross-spectral matrix. The non-parametric statistical testing at *p* < 0.01 was performed to evaluate the significant differences in global coherences at 40 Hz during the auditory stimulation conditions. The differences in coherences between task and baseline condition were quantified using coherence Z-statistic (Maris et al., 2007).

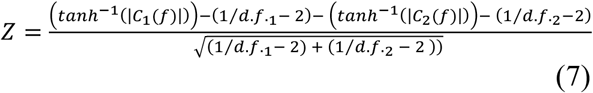

where *d. f*._1_ and *d. f*._2_ denote the degrees of freedom, *C*_1_(*f*) and *C*_2_(*f*) are coherence values during the first and the second conditions, respectively, at the frequency of interest (*f* = 40 Hz). The resulting Z-statistic was considered an observed test statistic, which was tested for significance after comparison with the null distribution as described above (see section 2.5.2).

#### 2.7.2. Directed sub-networks of ASSRs

After confirming the presence of a synchronised network, we calculated the multivariate Granger causality (GC) to establish the direction and strength of causal influence (Granger 1969). Subject-level nonparametric GC in the spectral domain were calculated among sources (Brovelli et al. 2004; Dhamala, Rangarajan, and Ding 2008). This involved nonparametric spectral matrix factorization (using Wilson’s algorithm) of the cross-spectral density to yield transfer function *H* and the noise covariance matrix ∑. Pairwise GC from Y to X at the frequency(*f*) can then be expressed as

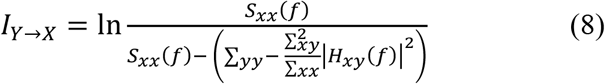

where *S_xx_*(*f*) is the total spectral power (auto-spectrum) and the denominator represents intrinsic power (total power minus the causal contribution) of the “effect” signal (Geweke 1982). Please note that since ASSR being phase-locked to the external signal, can result in spurious coherence estimation among sources. However, the calculation of GC reflects “true” connectivity, since, if the information content is same between X and Y, the GC value would be nearly zero. Particularly, as per eq (8) Y should contain additional information that is useful in predicting the future values of X, over and above that can be predicted from the past values of X alone.

Thereafter, measuring causality in both directions, one can locate the “source” and “effect” regions of the brain by comparing significant GC values in both directions. Since GC values follow unknown distribution, we adopt non-parametric statistical testing to assess significant GC spectral peaks among GC pairs (Brovelli et al., 2004). First, 1000 permuted data sets were generated by independently shuffling trials from each source pair. Shuffling trial order in this way abolished task-specific information while keeping the data pool same. Thereafter, GC was computed, and the maximum GC value was selected over the frequency range from each permuted data set (Ding, Chen, and Bressler 2006). Subsequently, a null distribution consisting of all GC values was constructed from the shuffled data set. GC peaks in unshuffled data were considered statistically significant when the observed GC value reached beyond the 99th quantile value (*p* = 0.01) of the null distribution. The multiple comparison problem was handled by Bonferroni corrections. We have also quantified the relative contribution of each source in the whole-brain causal network, as number of connections linked to each source. Mathematically, node degree was calculated as the number of edges connecting the source normalised by the maximum number causal connections each source could make i.e., (*N* – 1) *X* 2. In our case, we found 6 significant sources, so any source could maximally make 10 causal connections. Additionally, since our objective included identifying the role of trans-cortical communication in manifestation of right hemispheric dominance during different stimulation conditions, we specifically focused on the inter-hemispheric flow. Here, to maximize the number of measurements, Granger causality was calculated again from all voxels in the left PAC to all voxels in right STG during different stimulation conditions. Subsequently, we evaluated the effect on interhemispheric flow during different stimulation conditions by a non-parametric Kruskal-Wallis test, followed by Tukey-Kramer post-test.

## Results

### 2.8. Presence of ASSRs at sensor-level

In the time domain, evoked potential of mastoid channels showed steady-state activity in the time range of 250 – 1000 ms from the onset of periodic auditory stimuli (Figure 1A: Lower panel). In the frequency domain, we evaluated the power spectrum and strength of inter-trial phase clustering (ITPC) during auditory stimulation and baseline conditions across the frequency range of 3-46 Hz. We found significant ITPC during all auditory stimulation conditions (*p* < 0.01) only at 40 Hz (Figure 1B). Increase in the ITPC was a result of phase clustering around one region of polar space while the distribution of phase angles is uniform during the baseline condition (Figure 1C). Grand mean power spectra averaged over all electrodes and trials showed significant enhancement of spectral power at 40 Hz in binaural [*t* _(1970)_ = 4.19, *p* < 0.0001], monaural left [*t* _(1970)_ = 2.28, *p* < 0.0001] and monaural right condition [*t* _(1970)_ = 3.19, *p* < 0.0001] (Figure 1D).

### 2.9. Hemispheric asymmetry

Laterality indices were calculated to quantify the degree of asymmetry during ASSRs across different stimulation conditions at sensor level and source level (from bilateral Heschl’s gyri). At sensor level hemispheric LI were calculated for both 40 Hz spectral power and ITPC. Wherein, mean LI values were greater than zero during every auditory stimulation condition suggesting right hemispheric dominance ASSRs during binaural and both monaural conditions (Figure 2 B, Figure 2 C and Table S1). Particularly, for spectral power during the binaural condition, the group-mean hemispheric laterality index was LI = 0.023 and with 95 % lower and upper confidence interval was 95% CI = [0.010, 0.035]. During the monaural left condition (LI = 0.019, 95% CI = [0.007,0.032]) and monaural right condition (LI = 0.032, 95% CI = [0.019, 0.045]). One-way ANOVA among LI values across different stimulation conditions showed no significant difference in mean LI values across conditions (*F*(2,5910) = 1.01, *p* = 0.36). The laterality indices calculated from ITPC values also followed similar pattern. Specifically, during the binaural condition, (LI = 0.025, 95% CI = [0.014, 0.036]), monaural left condition (LI = 0.037, 95% CI = [0.025, 0.048]) and the monaural right condition (LI = 0.019, 95% CI = [0.008, 0.030]). Moreover, no main effect of auditory conditions was found on mean laterality indices (*F* (2, 5910) =2.51, p = 0.08).

**Figure 2.**
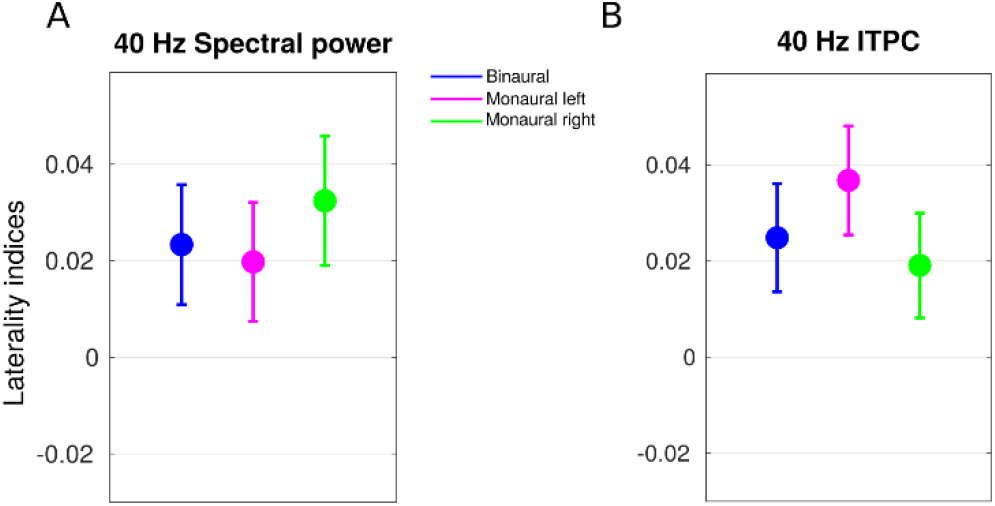
Hemispheric asymmetry: Group-level hemispheric laterality indices (LI) distribution for 40 Hz (A) spectral power and (B) ITPC during different stimulus conditions. The central node in each line represents mean of data while lower and upper boundary of the line represents the lower and upper limit of 95% confidence interval, respectively.

At source level LI were calculated for N100 response and ITPC from bilateral Heschl’s gyri to validate the presence of contralaterality effect during monaural condition (See SI). In line with earlier studies, clear contralateral dominance in early auditory potentials was observed for monaural conditions. Particular pairwise *p*-values that define effect of auditory conditions on the laterality values obtained from Tukey-Kramer post-test are listed in the Table S2.

### 2.10. Source-level functional organization of 40 Hz ASSRs

#### 2.10.1. Sources of ASSRs

Exact low-resolution brain electromagnetic tomography (eLORETA) was used to reconstruct whole-brain distribution of 40 Hz activity. The source activity was parcellated according to AAL atlas and tested for significance. The locations of significant sources during monaural left, monaural right and binaural conditions are shown in Figure 3. Anatomical labels corresponding to significant source and their respective t-statistic are summarised in Table 1. In total, six sources were found significant at p < 0.001 during every stimulation condition. Interestingly, locations of sources were same for binaural and both monaural conditions see Table 1), namely, bilateral Heschl’s gyri, bilateral precentral gyri, right superior temporal gyrus and right inferior frontal gyrus (triangular part). Hence, 4 sources were located in right hemisphere and 2 sources in left hemisphere for every stimulation condition. Though, the sources were same the source power of different regions differed across auditory stimulation conditions (Table 1).

**Figure 3.**
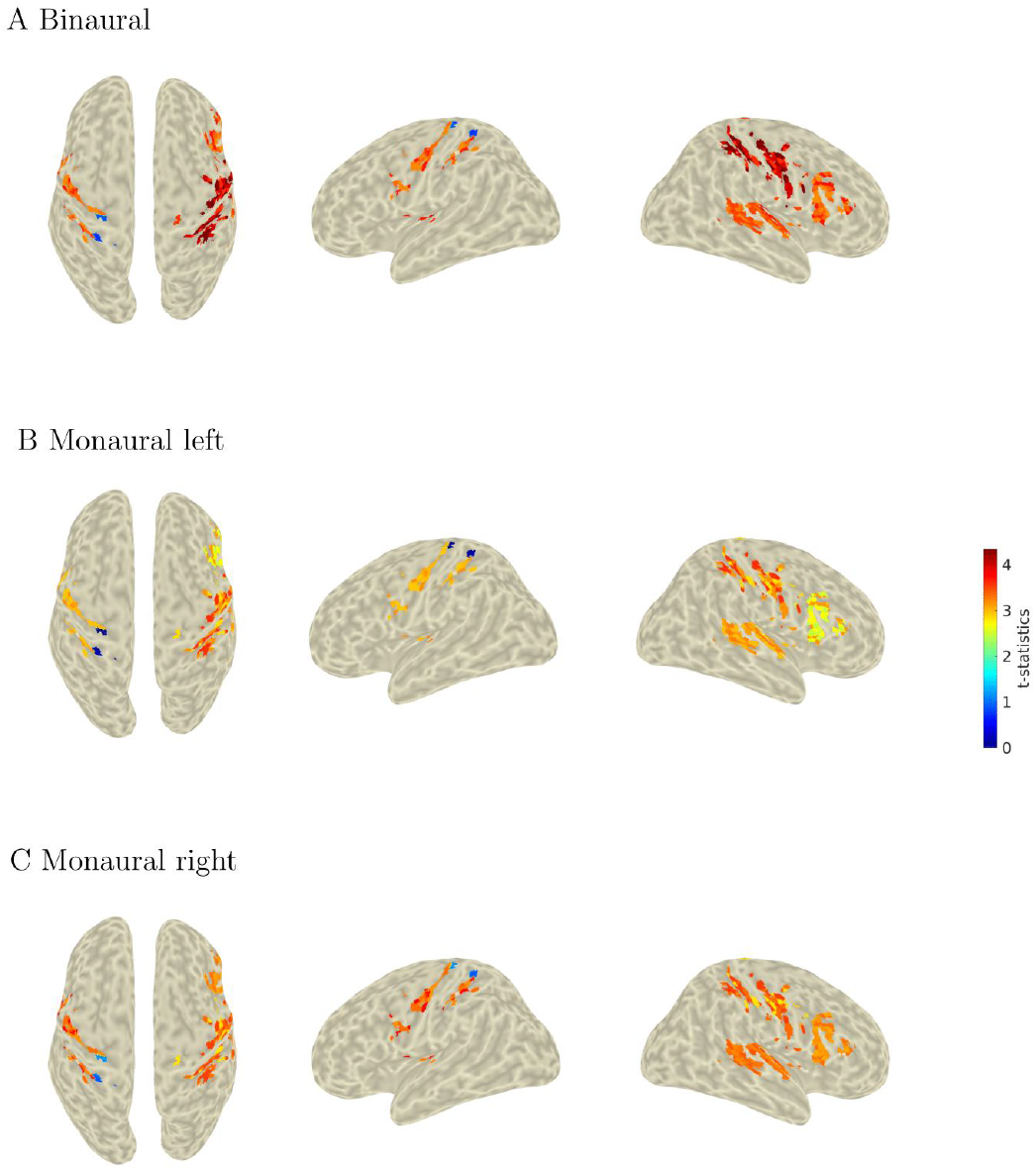
Sources of 40 Hz ASSRs: source power rendered over cortical surface derived from Colin27 brain.

**Table 1:** Anatomical labels (according to AAL parcellation) of 40 Hz ASSRs sources along with their corresponding power (t-statistic between auditory stimulation and baseline condition).

|  | Binaural | Monaural left | Monaural right |
| --- | --- | --- | --- |
| Right Heschl's gyrus | 3.81 | 3.25 | 3.46 |
| Left Heschl's gyrus | 3.69 | 3.24 | 3.28 |
| Right Superior temporal gyrus | 3.52 | 3.12 | 3.31 |
| Right Inferior frontal gyrus (triangular part) | 3.39 | 2.70 | 3.17 |
| Right Precentral gyrus | 4.01 | 3.17 | 3.34 |
| Left Precentral gyrus | 3.35 | 3.01 | 3.34 |

#### 2.10.2. Functional brain networks underlying ASSR

Global coherence measures the extent of coordinated neuronal activity over the whole brain (Cimenser et al., 2011; Kumar et al., 2016). The enhancement in global coherences among ASSR sources was evaluated by nonparametric statistical testing (Maris et al., 2007), employing Z-statistic as the test statistic. In line with our hypothesis, we observed a stimuli-specific enhancement of global coherence at 40 Hz, revealing a highly selective large-scale synchronization of neuronal assemblies at this frequency during binaural [*Z_20_* = 1.35, *p* < 0.0001], monaural left [*Z_20_* = 1.14, *p* < 0.0001], and monaural right [*Z_20_* = 1.17, *p* < 0.0001] conditions (Figure 4). The increase in whole-brain global coherence motivated for pairwise connectivity analysis employing Granger causality.

**Figure 4.**
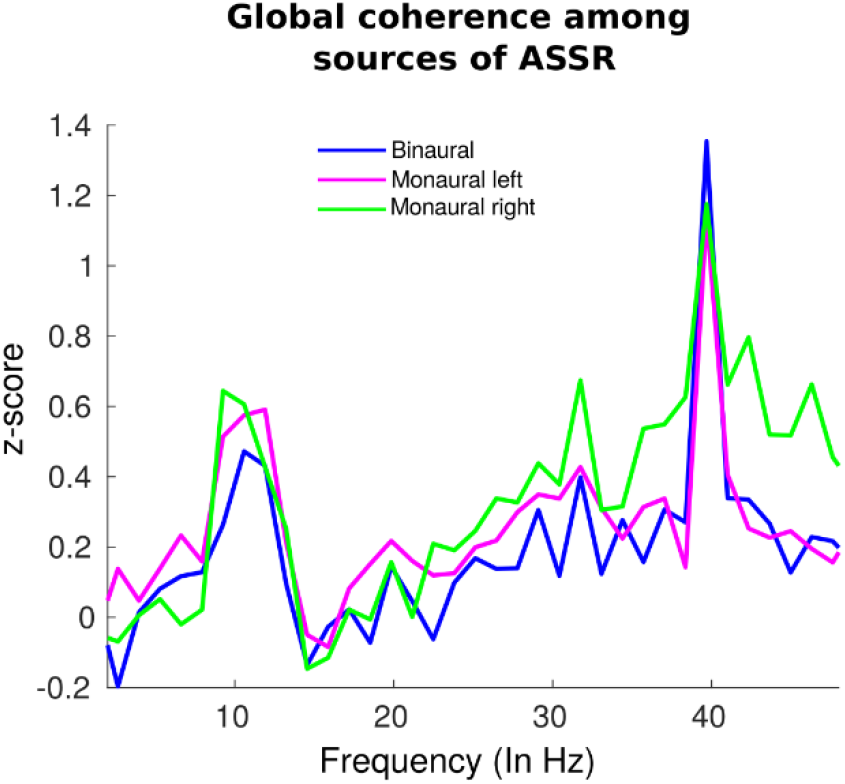
Presence of large-scale network among ASSRs: Enhancement in global coherence during auditory conditions was measured by z-statistic representing the normalized difference of coherence in task conditions relative the baseline condition.

Multivariate non-parametric spectral Granger’s causality (GC) was computed in all possible sources of ASSRs to measure source-level directed networks that may contribute to the whole-brain global coherence contribution (Geweke 1982; Granger 1969). Heatmaps in Figure 5 (left panel) show GC values among all possible source pairs. Significant Causal flow were depicted in the glass brain (Figure 5; Right panel) and also tabulated in Table 2 with corresponding GC values.

**Figure 5.**
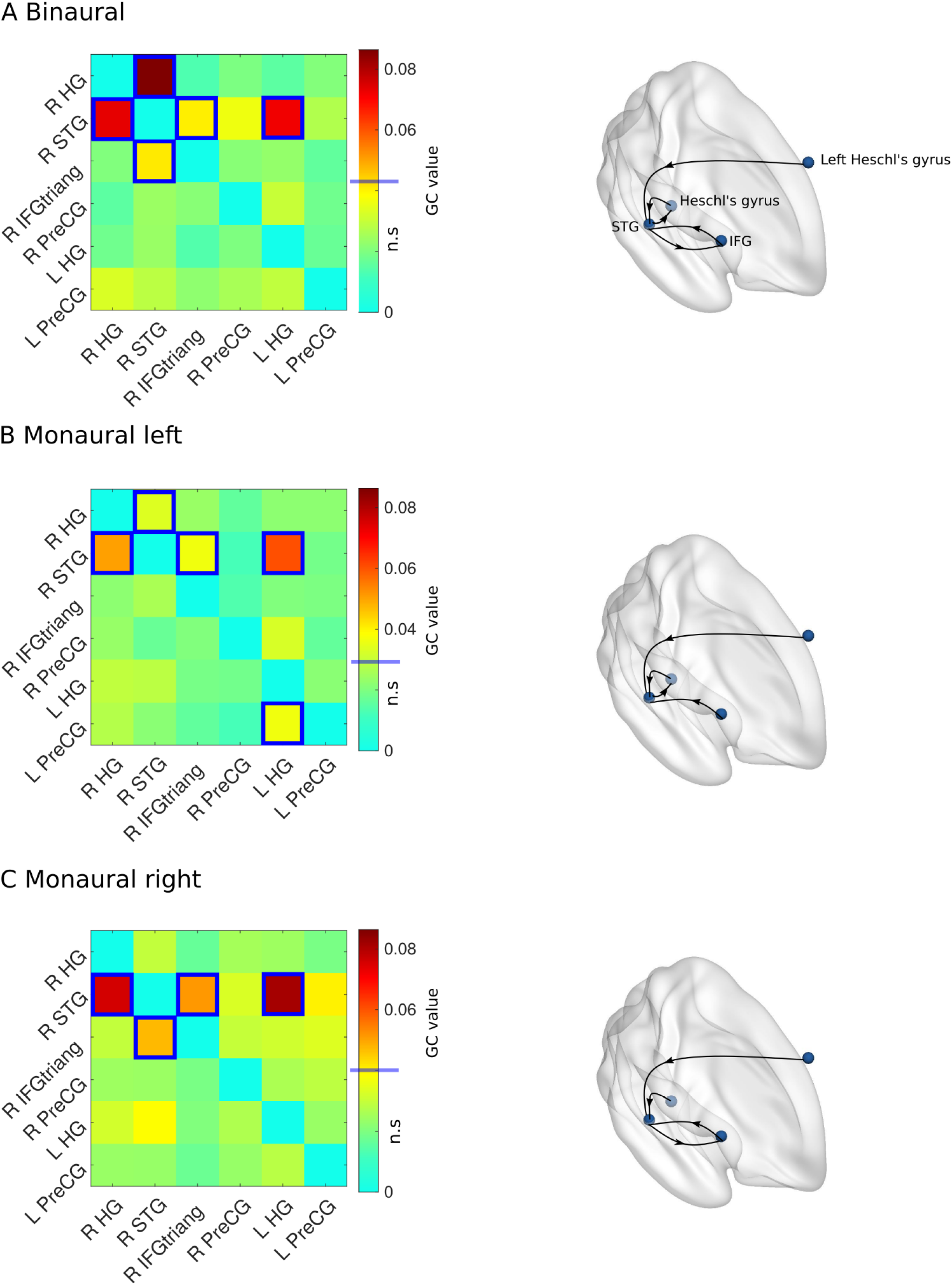
Directionality among recruited sub-networks: Left panel: heatmaps depicting pairwise GC values among ASSRs sources. 1st and 4th quadrant compose of GC pairs within right and left hemisphere, respectively. 2nd and 3rd quadrant depict respective interhemispheric GC flows. Significant GC values (p > 0.01) tested against surrogate data using a permutation-based non-parametric statistical test, were bordered by blue boxes. Schematic representation of frequency specific directed interactions that underlie 40 Hz ASSRs. Each black arrow represents a causal flow among brain regions denoted by blue nodes.

**Table 2:** Pairwise list of causally interacting sources pairs along with their respective causal strengths. Causal interactions among sources identified using Granger causality and significant causal interactions are illustrated in Figure 5.

| <u>Binaural</u> |  |  |
| --- | --- | --- |
| <u>From</u> | <u>To</u> | <u>GC value ($10^{-2}$)</u> |
| R STG | R HG | 8.6 |
| R HG | R STG | 7.3 |
| L HG | R STG | 7.2 |
| R STG | R IFGtriang | 4.1 |
| R IFGtriang | R STG | 4.1 |

| <u>Monaural left</u> |  |  |
| --- | --- | --- |
| L HG | R STG | 6.0 |
| R HG | R STG | 5.1 |
| L HG | L PreCG | 3.7 |
| R IFGtriang | R STG | 3.7 |
| R STG | R HG | 3.4 |

| <u>Monaural right</u> |  |  |
| --- | --- | --- |
| L HG | R STG | 8.2 |
| R HG | R STG | 7.6 |
| R IFGtriang | R STG | 5.1 |
| R STG | R IFGtriang | 4.7 |

Connectivity plots revealed the organization of directed network interaction during different auditory conditions (Figure 5; Right panel). Interestingly, at least three GC interactions were common in all three conditions; namely, 1) Left HG to Right STG, 2) Right HG to Right STG and, 3) Right STG to Right IFG. Specifically, outflows from bilateral HG reaching to the right STG, consequently involving unidirectional interhemispheric flow from left HG to right STG.

Presence of interhemispheric flow was in line with our hypothesis of involvement of trans-cortical flow during ASSR. During the monaural left and binaural conditions, there was bidirectional causal flow between right HG and right STG. During the binaural and monaural right condition, the right STG was bidirectionally connected to the right IFG. Node degree of ASSR sources (see Table 3) showed right STG was the most connected source among all ASSR sources having an average node degree of 0.43 followed by right HG (node degree = 0.17).

**Table 3:** Node degree of ASSRs sources in the whole-brain network of ASSRs during different auditory stimulation conditions. Fourth column show the average of node degree calculated as mean of node degree in all 3 conditions.

| Source Name | Binaural | Monaural left | Monaural right | Average |
| --- | --- | --- | --- | --- |
| Right Superior temporal gyrus | 0.50 | 0.40 | 0.40 | 0.43 |
| Right Heschl's gyrus | 0.20 | 0.20 | 0.10 | 0.17 |
| Right Inferior frontal gyrus | 0.20 | 0.10 | 0.20 | 0.17 |
| Left Heschl's gyrus | 0.10 | 0.20 | 0.10 | 0.13 |
| Left Precentral gyrus | 0.00 | 0.10 | 0.00 | 0.03 |
| Right Precentral gyrus | 0.00 | 0.00 | 0.00 | 0.00 |

Specifically, right STG was causally connected with bilateral HG and right IFG during every ASSR condition. Overall, three nodes from the right hemisphere (HG, STG and IFG) contribute to whole-brain causal network of ASSR while left hemisphere had two nodes (HG, Precentral Gyrus). For visualization purpose, we did not plot GC flow from left HG to left PCG during the monaural left condition.

Additionally, we found that the strength of inter-hemispheric flow was significantly different between at least two groups (χ (2,8097) =80.34, p < 0.0001). Particularly, both binaural and monaural right were significantly different from monaural left conditions (p < 0.0001) however there was no difference between binaural and monaural right conditions (p = 0.9). The difference in distributions of inter-hemispheric flow was visualised by ‘violin.m’ (Hoffmann, 2015). The maximum interhemispheric flow was present during the binaural condition, followed by monaural right and least during the monaural left condition (Figure 6).

**Figure 6.**
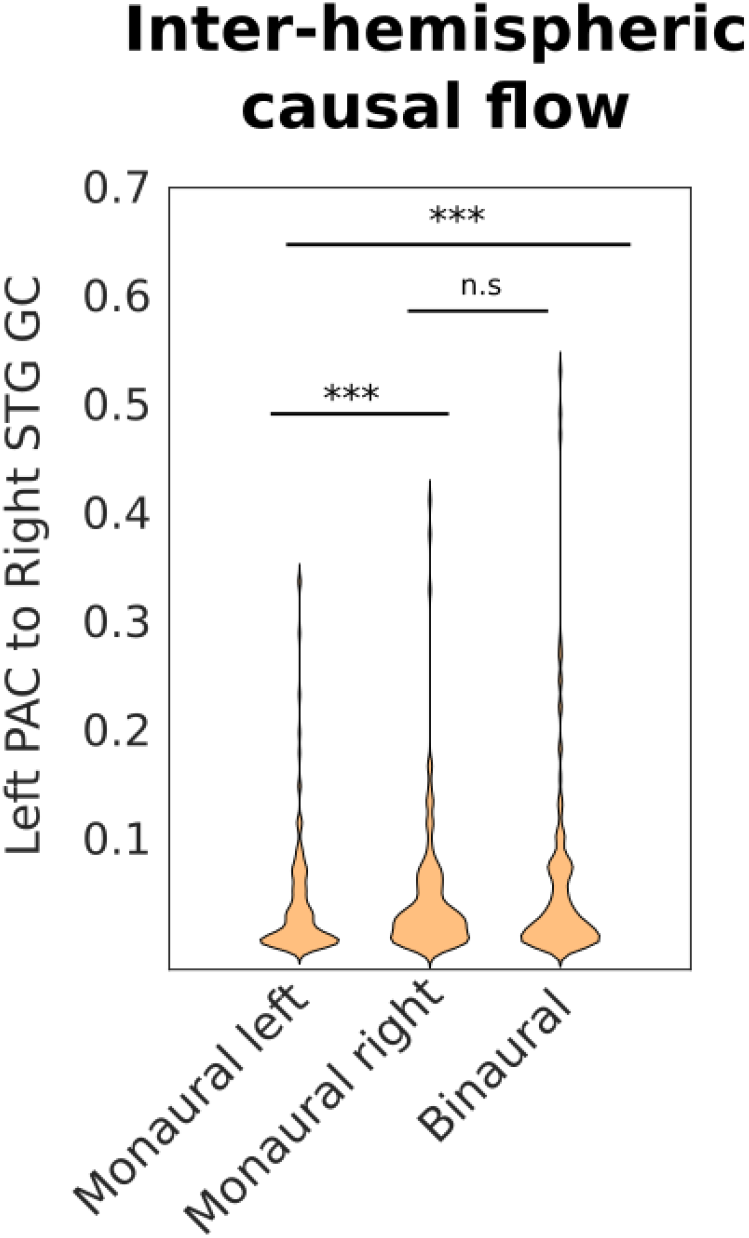
Interhemispheric causal flow: Variability in the GC strengths from left primary auditory cortex to the right STG during different auditory conditions.

## 3. Discussion

Our study attempted to reveal the directed interactions that unfold during the entrainment of periodic auditory stimuli of tonal nature in brain dynamics. We validated the presence of right hemispheric dominance of 40 Hz ASSR using spectral analysis and phase-locked components of the ASSR as estimated via Inter-trial phase clustering (ITPC). The main results were same in both measures as non-phase locked component in ASSRs is very less, as reported by our recent study (Singhal et al., 2023). Hence, spectral power is mostly attributed to the phase-locked component of ASSR. The stimuli-driven sources of neuronal oscillations were present across the hierarchy of cortical auditory pathways — Heschl’s gyrus (HG), superior temporal gyrus (STG), precentral gyri, inferior frontal areas corroborating with the previous fMRI, PET and EEG/MEG studies (Farahani et al., 2020; Poelmans et al., 2012; Reyes et al., 2004; Steinmann et al., 2011). Furthermore, we explored the information flow organization in these structures using effective connectivity analysis. We found that the right STG serves as an integrative area, receiving inputs from both primary auditory cortices, wherein information from the left primary auditory cortex is received via trans-cortical pathways, the organizational symmetry of which was dependent on the ears being stimulated. Furthermore, our directional network analysis suggests bidirectional interactions between right STG and frontal regions may provide efficient information exchange required for a comprehensive mapping of the auditory environment that ultimately manifests as right hemispheric dominance. The present findings may provide not only a network-level basis for right hemispheric dominance during 40 Hz ASSRs but also a clue about how the variability in the strength inter-hemispheric flow is dependent on the ear being stimulated. For instance, due to the pre-existing contralateral dominance, right PAC receives least auditory input thereby require greater inter-hemispheric flow to compensate for the asymmetric input.

### 3.1. Sources of ASSR

A significant oscillatory response was seen in bilateral HG during every condition. HG residing in the primary auditory cortex is known to be the first cortical structure that receives auditory inputs (Hackett, 2015). Both monaural conditions showed contralateral dominance in early evoked potentials (See SI). Contralateral dominance of primary auditory areas during monaural stimulations is a well-established phenomenon reported in several fMRI, PET and EEG/MEG studies and attributed to the crossing of ascending anatomical fibres at the brainstem level (Hackett, 2015; Kaas & Hackett, 2000; Langers et al., 2005). In addition to the classic auditory pathway, we also report activation beyond the primary auditory cortex, for instance, bilateral pre-central gyri, which is in line with earlier findings on reconstructed 40 Hz ASSRs sources with equivalent dipole modelling (Farahani et al., 2020). Interestingly, significant activation in the right inferior frontal gyrus (triangular part) (rIFG) was also found during every condition of ASSR. Right IFG, an analogue of Broca’s area in the right hemisphere has been suggested to process sound length and attending to pitch/rhythms (Plakke & Romanski, 2014; Wang et al. 2015). Though most sources of 40 Hz ASSR are well established, the enhancement of global coherence among sources in the present study reveals inter-areal brain synchronization among sources, a prerequisite of communication among neuronal assemblies (Bressler & Menon, 2010; Fries, 2005). Further analysis of directed information flow among sources provides a much clearer understanding of the role of observed sources.

### 3.2. Interhemispheric transfer

During every stimulation condition, there were causal flows originating from both primary auditory cortices reaching the right STG, consequently requiring an interhemispheric causal flow from the left PAC (Figure 5). There are anatomical fibres that support this information flow. In general, interhemispheric transfer of information is achieved via the corpus callosum (Andoh et al., 2015). The corpus callosum provides a functionally relevant scaffold for mediating proper communication across both hemispheres (Zaidel & Iacoboni, 2003). Although the inter-hemispheric flow was present during every auditory stimulation condition there was differences in strength of interhemispheric flow during different conditions (Figure 6). The observed differences in the strength of interhemispheric pathway during different task conditions suggest its differential functional requirements. Due to existing afferent structural constraints, irrespective of the ear being stimulated both PAC receives some amount of acoustic information in tandem with the manifestation of contralateral dominance during monaural conditions (Langers et al., 2005). The right hemispheric specialization to process rhythmic features of acoustic input would require transfer of information from left PAC to specialised centres present in the right hemispheres. Hence, the monaural right condition would naturally require higher interhemispheric flow to compensate for the early asymmetric inputs. Alternatively, during the monaural left condition, wherein the information is already dominant in left hemisphere (Figure S1) would show least involvement of inter-hemispheric flow. The strength of inter-hemispheric flow was maximum during the binaural condition (Figure 6). In general, binaural condition requires more trans-cortical communication to integrate incoming bilateral acoustic inputs in addition to the transfer of primary information from PAC to the secondary auditory cortex. The complexity in dissociation of these two kinds of information prevents a straightforward explanation of inter-hemispheric flow during binaural conditions.

### 3.3. Integrative nature of STG

After initial processing in bilateral HG, information related to ASSR is feedforwarded to the right superior temporal gyrus (rSTG). Several studies of ASSRs have reported prominent activations in rSTG (Edgar et al., 2016; Kim et al., 2019; Mäkelä and Hari 1987). We observed rSTG showing maximum inward causal information flow in the whole network (See Figure 5 and Table 2). This type of causal information flow from primary sensory areas to intermediate areas of the auditory pathway suggests the integrative nature of rSTG mediating acoustic-pattern analysis. Ross and colleagues proposed that the right auditory cortex process temporal regularities associated with pitch processing of incoming sound (Ross et al. 2005). In the context of the earlier view, rSTG can be considered as a specialised region for detecting regularities in acoustic input and subsequently sending information to “higher-order” frontal cortices. Also, our analyses revealed that rSTG is bidirectionally connected to the right inferior frontal gyrus (BA45). STG to frontal gyrus causal flow can be considered a bottom-up process, while causal influence from frontal to right STG can be seen as a top-down modulation or relevant for predicting incoming sensory inputs. Right IFG (rIFG) areas are known to be associated with identifying features like sound length and attending to pitch/rhythms plausibly utilising periodic cues received from STG (Plakke & Romanski, 2014). Activity in rIFG along with activation in bilateral primary auditory cortex was also reported by Reyes and colleagues using PET imaging in response to 40 Hz amplitude modulated (AM) tone (Reyes et al. 2005). Several studies have reported anatomical projections from STG to the frontal cortex (Hackett 2011; Kaas and Hackett 2000; Plakke and Romanski 2014). Wang and colleagues identified a functional network comprising the frontal cortex and superior temporal regions that are sensitive to tone repetition patterns, which is associated with human’s unique ability for language processing (Wang et al. 2015). Here we show, this fronto-temporal network that is central to processing of periodic structure of auditory stimuli, is mediated by bidirectional interactions between rSTG and right frontal regions.

### 3.4. Right hemispheric dominance

The present findings could give us a possible basis for right hemispheric dominance during 40 Hz ASSRs, as observed in several past studies by exploring the directed information flow among cortical sources. We argue that the right hemispheric lateralization of 40 Hz can be attributed to the right STG and its bidirectional interaction with the right frontal region. Furthermore, the activation of the pattern was similar in every condition, suggesting it as a central network of processing of periodic auditory stimuli and facilitator of brain-wide entrainment of cortical rhythms. This is in line with the previous finding that suggested tonal or melodic stimuli are predominantly processed in the right hemisphere while speech and language stimuli showed left hemisphere dominance (Albouy et al., 2020; Zatorre and Belin, 2001; Ross et al., 2005; Zatorre and Gandour 2008) thus, following the same reverberating theme of right hemispheric dominance in music processing.

## 4. Methodological considerations

We have employed distributed source modelling based on individual-subject anatomy to account for variability in participant head size. Our source analysis was not limited to cortical regions, but we did not detect any significant activations in the subcortical areas of the brain. However, it should not be assumed that subcortical sources do not play a role in the whole-brain processing of the ASSR. There are substantial studies about the role of thalamocortical circuits in mediating both auditory-cortices and generation of ASSR (Lee, 2013; Li et al., 2018; Steinmann et al., 2011). However, in our study, the detectability of thalamus and other sub-cortical regions was limited by several methodological factors. In principle, sub-cortical regions of the brain are prone to signal cancellation due to their deep location and irregular cell architecture, causing cortical activity to dominate over sub-cortical activity (Attal et al., 2007; Hnazaee et al., 2020). Consequently, poor signal-to-noise ratio for deeper sources (Halder et al., 2019), particularly in estimating the 40 Hz ASSR as higher frequency phases are more susceptible to distortion during propagation to the scalp. Emerging studies suggest the possibility of detecting sub-cortical sources by limiting the source analysis to pre-defined areas or by defining the region of interest larger than its actual volume (Coffey et al., 2016; Farahani et al., 2021). However, in our study, both of these approaches would be counter-intuitive as we did not have prior assumptions about the locations of source-level network dynamics. The influence of sub-cortical activity on whole-brain network auditory processing could be potentially delineated in a future EEG-fMRI study where source network locations can be identified over long-time scales from BOLD activity and source-level EEG time series reconstructed from a spatial-filter that is better receptive to thalamic signals.

## 5. Conclusion

Comprehensive source-level network analysis provides a plausible explanation of hemispheric asymmetry of 40 Hz ASSR during binaural and both monaural conditions. Right hemispheric dominance emerges due to bidirectional flow between the right STG and right frontal area. Inter-hemispheric flow compensates for pre-existing contralateral dominance in early sensory processing to yield right hemispheric specialization of tonal processes. Overall, while the source locations and direction of information flow were similar across both monaural and binaural conditions, their differential strength of activation underlies the processing of different auditory environments.

## Acknowledgements

The study was supported by NBRC Core funds and by grants Ramalingaswami fellowship, (BT/RLF/Re-entry/31/2011) and Innovative Young Bio-technologist Award (IYBA), (BT/07/IYBA/2013) from the Department of Biotechnology (DBT), Ministry of Science and Technology, Government of India to AB. AB also acknowledges the support of Centre of Excellence in Epilepsy and MEG (BT/01/COE/09/08/2011) from DBT. DR was supported by the Ramalingaswami fellowship (BT/RLF/Re-entry/07/2014) from DBT. We sincerely thank late Dr. Jeffrey Michael Valla for his constructive comments on an earlier version of this manuscript.

## Supporting Information

### SI Methods

Early auditory potentials were reconstructed employing time domain exact low-resolution brain electromagnetic tomography (eLORETA) (Pascual-Marqui, 2007). Similar to frequency domain eLORETA as described in the main manuscript (See Methods 2.6) we have utilised individual anatomy to reconstruct source waveforms while restricting the analysis to bilateral primary auditory cortices. Wherein, the respective coordinates of sources were obtained for both Heschl’s gyri based on the Automated anatomical labelling (AAL) atlas (Rolls et al., 2015) (See figure S1 A; left panel). Sensor-level covariance matrices were computed for each condition from the 20 ms time series centred around N100 response. Remaining procedure kept same as frequency domain source analysis (See Methods 2.6) to obtain subject and trial wise source waveforms for each condition. We also calculated ITPC from same areas based on the Fourier coefficients obtained from frequency domain eLORETA (See methods 2.7). Thereafter, laterality indices were computed for both N100 amplitude and ITPC between bilateral auditory cortices followed by statistical testing of asymmetry in individual condition and difference between each condition as per methods 2.5.3.

**Figure S1:**
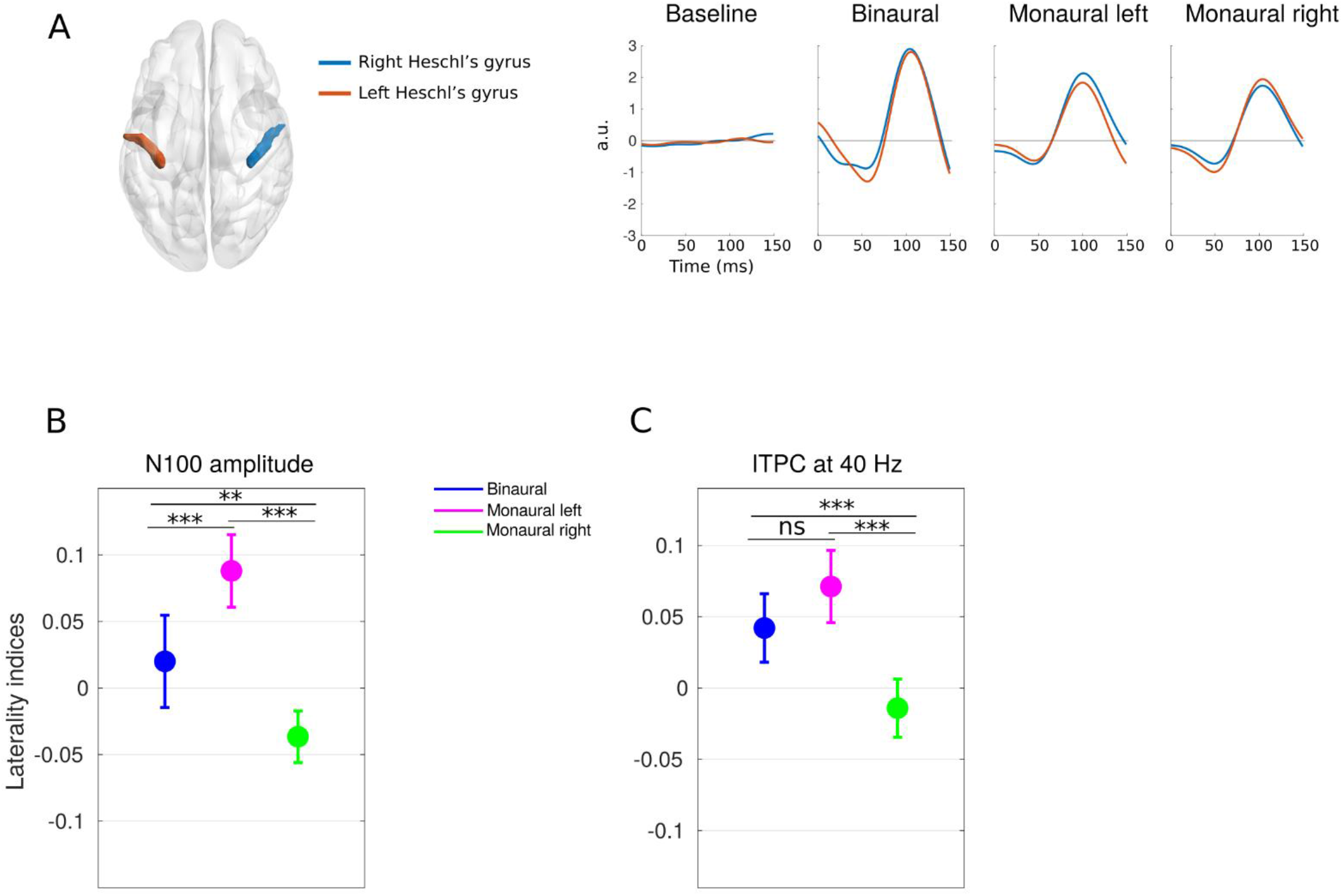
Asymmetry at the primary auditory cortices: A) Location of left and right Heschl’s gyri (left panel) and Grand average source waveforms (Right panel). B and C Mean and 95 % confidence interval of laterality indices of (B) N100 amplitude and (C) ITPC from Heschl’s gyri.

### SI Results

Source wave forms were low-pass filtered at 20 Hz to avoid distortion of ERP from high frequency activity. The source waveform analysis revealed a clear N100 response in both Heschl’s gyri across all ASSR conditions. There were significant amplitude differences observed between the left and right Heschl’s gyri, consistent with previous studies. Asymmetry in the responses were quantified using LI analysis. Particularly, binaural condition showed bilateral response while both monaural conditions were clearly contralaterally dominant. Particularly, during binaural condition the group-mean laterality index was LI = 0.020 and with 95 % lower and upper confidence interval was 95% CI = [-0.015, 0.055]. During the monaural left condition (LI = 0.088, 95% CI = [0.061,0.115]) and monaural right condition (LI = −0.037, 95% CI = [−0.056, −0.017]). One-way ANOVA among LI values across different stimulation conditions showed significant difference in mean LI values across conditions (F (2,57) =22.03, p < 0.001).

Right hemispheric dominance was observed for binaural condition showing group-mean laterality index was LI = 0.042 and with 95 % lower and upper confidence interval was 95% CI = [0.018, 0.066]. Though there was significant difference between monaural left and monaural right condition (p < 0.001), clear contralateral dominance was only present in monaural left condition i.e., (LI = 0.071, 95% CI = [0.046,0.097]). Monaural right response also not clearly left lateralised having mean LI = −0.014, and 95% (CI = [−0.034, 0.006]). Overall, there was significant difference in mean LI values across conditions (F (2,57) =15.12, p < 0.001).

### SI Discussion

There is substantial evidence that contralateral projection of ascending auditory pathway leads to contralateral representation in the primary auditory cortices, primarily involved in early auditory processing (Hackett, 2011; Hackett, 2015). In line with these findings, we observed contralateral dominance of N100 and ITPC measures at primary auditory cortices during monaural conditions (see Figure S1 and Table S1). Essentially, both measures are associated with distinct aspect of auditory processing. For instance, The N100 measure, which is an early auditory potential, showed clear asymmetry and varied among different auditory conditions (see Table S1 and S2). On the other hand, an increase in the ITPC is associated with late auditory potentials that are right hemispheric dominant for 40 Hz ASSR.

**Table S1:**
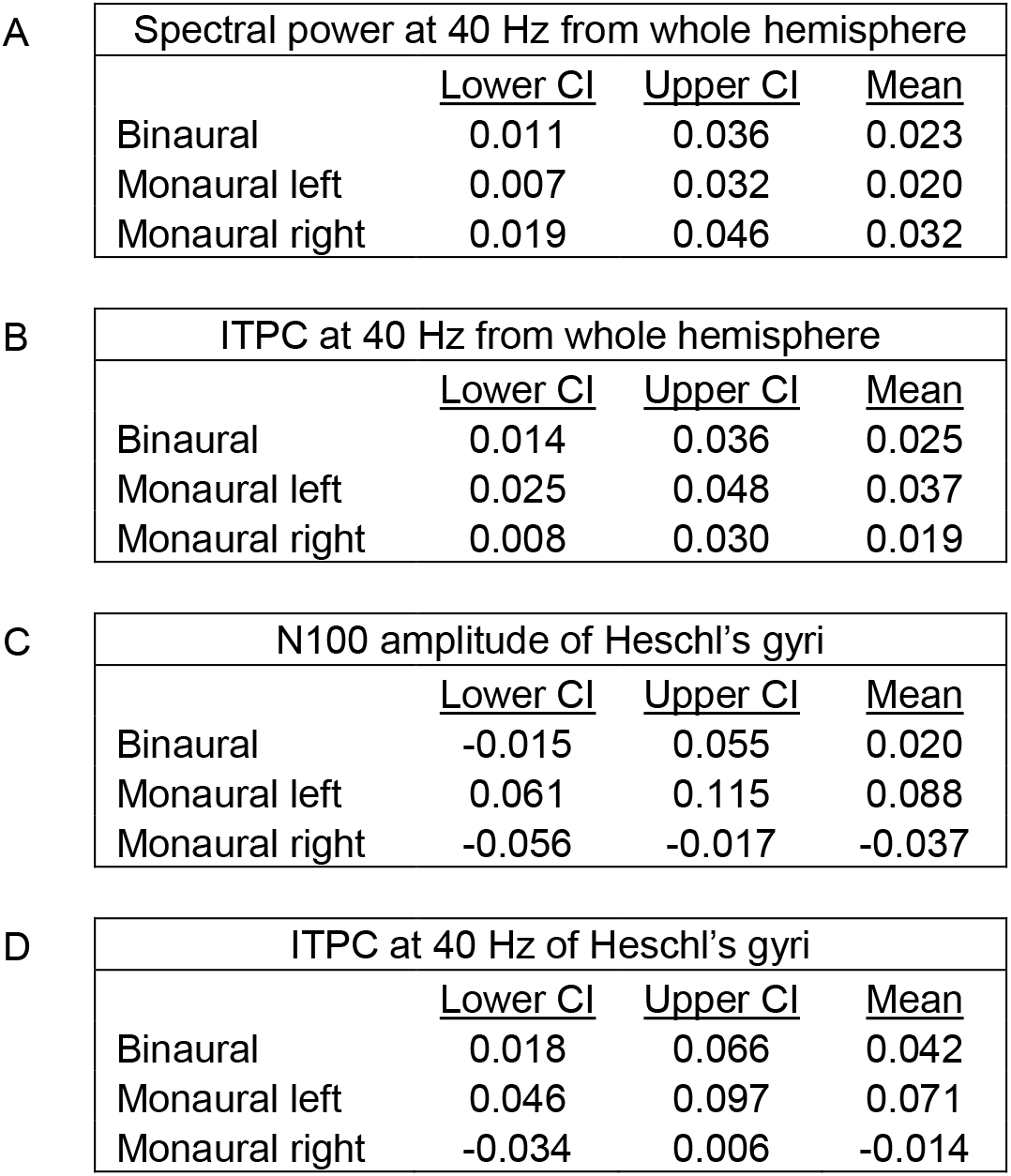
Mean and 95 % confidence interval (CI) of laterality indices during different auditory conditions. (A) and (B) measured at sensor level (See methods 2.5.3). (C) and (D) measured at source level (Heschl’s gyri). (C) N100 amplitudes were calculated from source waveforms derived from time domain eLORETA (See supplementary methods) and (D) ITPC were calculated from Fourier transforms obtained from frequency domain eLORETA (See methods 2.7).

**Table S2:**
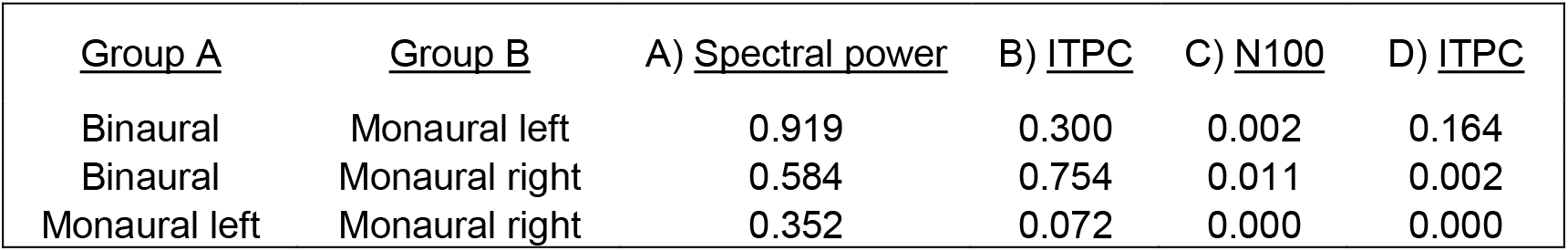
Pairwise p-values after multiple comparisons test Mean and 95 % confidence interval (CI) of laterality indices during different auditory conditions. (A) and (B) measured at sensor level (See methods 2.5.3). (C) and (D) measured at source level (Heschl’s gyri). (C) N100 amplitudes were calculated from source waveforms derived from time domain eLORETA (See supplementary methods) while (D) ITPC were calculated from Fourier transforms obtained from frequency domain eLORETA (See methods 2.7).

## Footnotes

1 A subset of this data (10 volunteers) were used in a Methods paper Halder, T; Talwar, S., Jaiswal, A.K., Banerjee, A.(2019): Quantitative evaluation in estimating sources underlying brain oscillations using current source density methods and beamformer approaches. eNeuro. 2019 Jul-Aug; 6(4): ENEURO.0170-19.2019.

